# Histone code dictates fate biasing of neural crest cells to melanocyte lineage

**DOI:** 10.1101/702332

**Authors:** Desingu Ayyappa Raja, Yogaspoorthi Subramaniam, Vishvabandhu Gotherwal, Jyoti Tanwar, Rajender Motiani, Sridhar Sivasubbu, Rajesh S Gokhale, Vivek T Natarajan

## Abstract

In the neural crest lineage, progressive fate-restriction and stem cell assignment are critical for both development and regeneration. While the fate-commitment events have distinct transcriptional footprints, fate-biasing is often transitory and metastable, and is thought to be moulded by epigenetic programs. Hence molecular basis of specification is difficult to define. In this study, we establish a role of a histone variant *H2a.z.2* in specification of melanocyte lineage from multipotent neural crest cells. Silencing of *H2a.z.2* reduces the number of melanocyte precursors in developing zebrafish embryos, and from mouse embryonic stem cells *in vitro*. We demonstrate that this histone variant occupies nucleosomes in the promoter of key melanocyte determinant *Mitf*, and enhances its induction. CRISPR-Cas9 based targeted mutagenesis of this gene in zebrafish drastically reduces adult melanocytes, as well as their regeneration. Thereby our study establishes a histone based specification code upstream to the core gene regulatory network in the neural crest lineage of melanocytes. This epigenetic code renders a poised state to the promoter of key determinant and enhances activation by external instructive signals thereby establishing melanocyte fate identity.

## Introduction

Cell fate determination is driven by a cascade of transcription factors that are in-turn governed by external cues (Kawakami and Fisher, 2011; Martik and Bronner, 2017). The multipotent neural crest sets a paradigm for stem cell pluripotency and lineage specification, and serves as a model to understand fate decisions. Recent data emerging from single-cell transcriptomics of neural crest cells (NCCs) demonstrates emergence of fate bias or specification that further unfolds to culminate in fate commitment (Soldatov et al., 2019) or determination. NCC’s and NCC-derived progenitor population give rise to melanocytes during embryonic development, and adult skin homeostasis and regeneration respectively (White and Zon, 2008). Ability to trace cells visually, combined with the extensive literature on gene regulatory network, render melanocytes as a preferred choice to trace mechanisms behind fate decisions.

Almost all aspects of melanocyte biology, from lineage commitment to differentiation, is controlled by the central transcription factor MITF (Goding and Meyskens, 2006; Kawakami and Fisher, 2017). WNT and BMP signalling (Jin et al., 2001; Takeda et al., 2000), and downstream transcription factors SOX10 (Marathe et al., 2017), PAX3 (Watanabe et al., 1998), FOXD3 (Kos et al., 2001) and LEF1 (Dunn et al., 2000), form the core of gene regulatory network that control *Mitf* induction and mediate melanocyte specification and determination. While the upstream instructive signals and downstream orchestrators are well appreciated, cell intrinsic modulators that mediate specification events remain obscure. Epigenetic mechanisms, by altering chromatin accessibility, could provide an intervening regulatory layer and are likely to play a key role in determining the fate of NCC lineages.

Epigenetic regulators including histone deacetylase HDAC1 (Ignatius et al., 2008; Ignatius et al., 2013) involved in locus specific histone deacetylation, DNA methyl transferase DNMT3b, that causes base methylation (Rai et al., 2010), and histone variant H3.3 that replaces the core histone H3, are involved in the derivation of specific cell types from NCCs (Cox et al., 2012). Another histone variant H2A.Z, encoded by two genes *H2a.z.1* and *H2a.z.2*, has been implicated in NCC differentiation (Punzeler et al., 2017; Sivasubbu et al., 2006). Recent studies demonstrate H2A.Z to play a crucial role in human metastatic melanoma survival and proliferation (Vardabasso et al., 2015) and involvement of H2A.Z chaperone PWWP2A in NCC differentiation (Punzeler et al., 2017). Since melanoma recapitulates neural crest fate, (Kaufman et al., 2016), it is possible that histone variants dictate the fate of melanocytes from neural crest derived progenitors.

In this study, using zebrafish as well as mouse embryonic stem cell-derived melanocyte models we demonstrate a selective role for H2A.Z.2 in fate-biasing of NCCs to melanocytes. Further, we show that this variant occupies the promoter of *Mitf* and upstream regulator *Sox9*, and governs the inducibility of *Mitf* under appropriate cues. A series of CRISPR targeted mutations in *h2a.z.2* reinforce its central role during pigmentation and the related process of melanocyte regeneration in adult zebrafish. Thereby, we establish a histone code upstream to the transcription factor network that enables cell-intrinsic responsiveness to external instructive signals. Our study adds a new dimension to the recent developments in tracing the lineage map of NCC derivatives and provides a molecular basis to the fatebias observed during specification.

## Results

### Histone variant *h2a.z.2* but not *h2a.z.1* is involved in the derivation of melanocytes in zebrafish

Across vertebrates, the histone variant H2A.Z is encoded by two distinct paralogs H2A.Z.1 (Z1) and H2A.Z.2 (Z2), that are 97% identical at the protein level, and hence difficult to distinguish experimentally (Dryhurst et al., 2009; Matsuda et al., 2010). Disruption of Z1 is not compensated by intact Z2 and the knockout mouse embryos fail to survive (Faast et al., 2001). Therefore, to decipher their individual contribution to the differentiation of neural crest cells (NCCs), especially to melanocytes, we adopted a morpholino based transient knockdown strategy in developing zebrafish embryos. Upon knock down of Z1, gross embryonic deformities and poor development were observed, as has also been reported previously (Madakashira et al., 2017; Murphy et al., 2018; Sivasubbu et al., 2006). Z2 morphants only had milder effects on gross morphology, and they presented a drastic reduction in pigmentation (Fig 1A-D).

**Figure 1:**
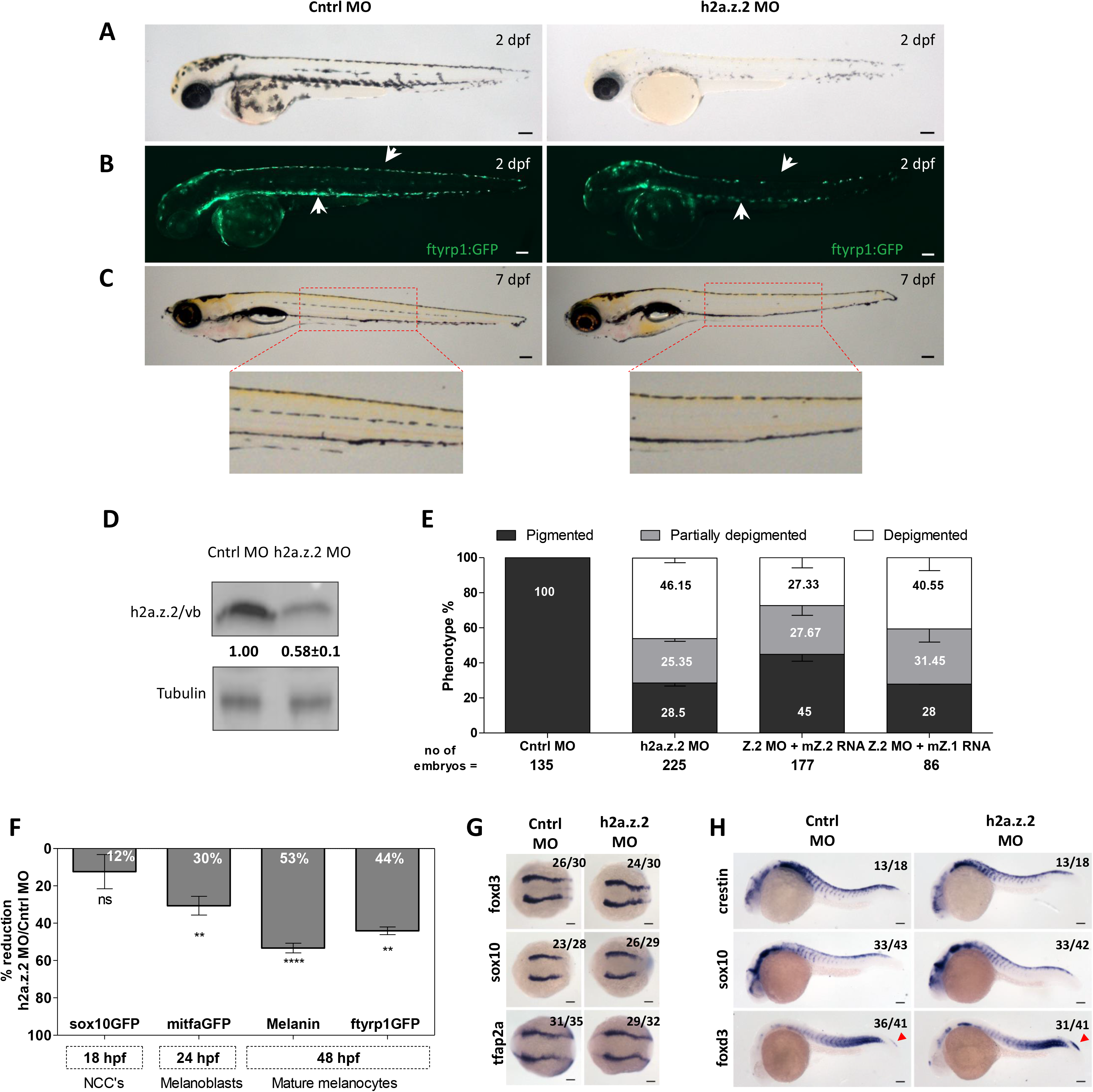
*H2a.z.2* controls the melanocyte numbers during zebrafish development. (A) Bright field images of control morpholino (control MO) and H2a.z.2 morpholino injected embryos (Z2 MO) at 2 days post fertilization (dpf). (B) Fluorescence image of *Tg(ftyrp1:GFP)* that tags differentiating melanophores, in control and Z2 MO embryos at 2dpf. (C) Bright field images of control and Z2 MO at 7 dpf. (inset) Enlarged view of the lateral line melanophores. (D) Western blot analysis of control and Z2 MO embyos at 2 dpf, carried out using an antibody that recognizes both H2A.Z.1 and H2A.Z.2 proteins. (E) Bar graphs represent means±SEM of percent embryos with varying degree of pigmentation (depigmented, partially pigmented and normally pigmented) scored manually at 2dpf, from embryos injected with control MO, Z2 MO, Z2 MO along with mouse *H2a.z.1* mRNA or mouse *H2a.z.2* mRNA. (F) Inverted bar graphs represent mean ± SEM (n=3) in the percent reduction in cell numbers of Z2 MO compared to control in various marker lines. Time of assessment of labelled cells, their identity and the transgenic line used are indicated. (G) Whole mount RNA in situ hybridization (WISH) based expression pattern of early neural crest markers *foxd3*, *sox10* and *tfap2a* at 11hpf. (H) WISH based expression pattern of neural crest markers *crestin*, *sox10* and *foxd3* at 24hpf. Numbers in WISH images indicate frequency of the represented phenotype in total number of embryos analyzed. Scale bars 100 μm.

Pigmentation phenotype of Z2 MO could be recapitulated using splice block MO targeted to Z2, and also upon co-injection of Z2 MO with p53 MO, reiterating the specific nature of the phenotype (Supplementary Fig 1). Upon co-injection of Z2 MO along with mouse H2a.z.1 or H2a.z.2 mRNAs, a rescue of around 50% was observed only for Z2 mRNA (Fig 1E). Thereby demonstrating that Z2 has a distinct pigmentation role, and is not compensated by increased dosage of Z1.

### *h2a.z.2* depletion selectively reduces melanocyte numbers

The characteristic reduction in pigmentation of Z2 morphants could be either due to decreased pigment production or due to the absence of mature pigmented cells. At 2 days post fertilization (2 dpf), we observed a severe reduction in the number of ftyrp1:GFP +ve cells in Z2 morphants, notably in dorsal as well as the ventral pool of melanocytes above the yolk sac, clearly indicating that the number of melanocytes is affected (Fig 1B). Additionally at 6 dpf, the lateral line melanocytes were reduced in Z2 morphants, suggesting that the progenitor pool may be affected (Fig 1D). We therefore used various transgenic reporter lines from neural crest to melanocyte lineage to estimate respective cell numbers at different stages of development as indicated in (Fig 1F). We observed that Z2 morphants displayed approximately 45-50% reduction in the differentiating/differentiated melanophores and a 30% reduction in melanoblast numbers (Fig 1F). However, the sox10 +ve progenitor population remained unaffected in Z2 morphants. *h2a.z.2* is therefore likely to play a key role in the lineage commitment of melanocytes from neural crest derived progenitors.

### *h2a.z.2* dictates melanocyte and glial footprint in NCCs

Multipotent NCCs progressively undergo the process of fate specification and determination to give rise to melanocytes, craniofacial cartilage, adipocytes, smooth muscle cells, schwann cells and neurons (Donoghue et al., 2008; Le Douarin et al., 2004). Based on Whole mount In Situ Hybridization (WISH), we observed that the early neural crest gene expression of *foxd3*, *sox10* and *tfap2a* were not decreased in Z2 morphants (Fig 1G&H). This indicates that depletion of *h2a.z.2* could selectively affect a subpopulation of NCC derived progenitors from which melanocytes and possibly a few other cell types would be derived. Hence we resorted to identify gene expression changes in this seemingly homogeneous population for alterations that could explain decreased melanocyte numbers.

We created Z2 morphants in *Tg(sox10:GFP)* line and sorted the GFP +ve cells at 16-18 hpf. At this time point, neural crest derived *sox10* expression is retained and also lineage specific differentiation markers begin to emerge and the cell fate identity is established (Curran et al., 2010; Wagner et al., 2018). Microarray and pathway enrichment analysis of the GFP +ve cells revealed that the set of downregulated genes mapped to developmental pigmentation related processes (Fig 2A-C). Targeted analysis of the data indicated reduced expression of transcription factors (TFs) with known role in melanocyte lineage, such as *mitfa*, *tfap2a*, *tfap2e* and *sox9b,* as well as melanocyte migration and survival factors such as *kita* and *ednrb1a*. Similarly, we also observed decreased expression of glial lineage transcription factors such as *nfatc4*, *pax7*, *lmo4* etc. However, the neuronal lineage TFs such as *foxo1*, *foxo3* and *sox8* were upregulated in the Z2 morphants (Fig 2D). As these are gene expression changes in the isolated sox10 +ve cells, we interpret the decreased expression to indicate alterations in the subpopulations of differentiating cells derived from the NCCs.

**Figure 2:**
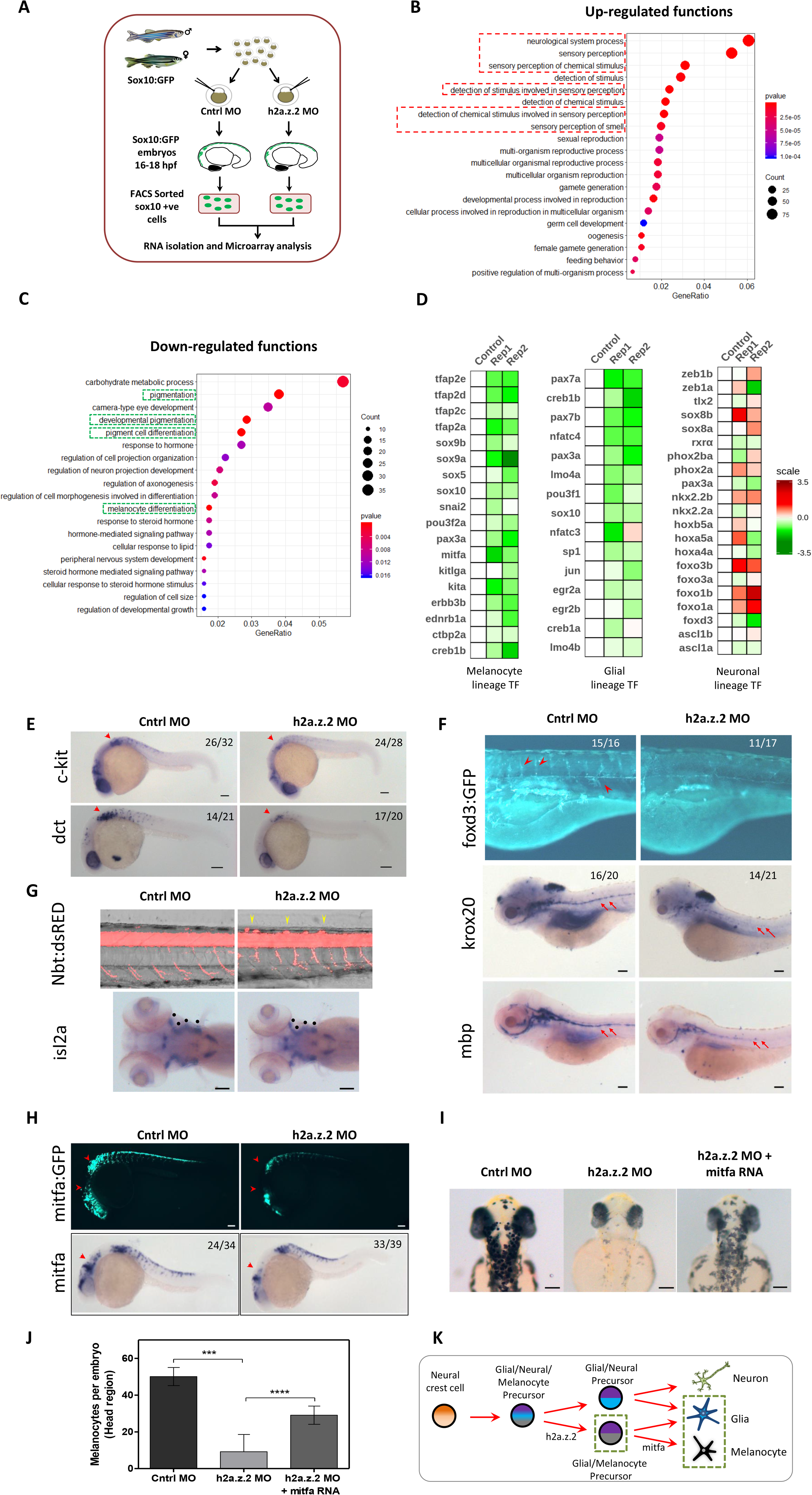
*H2a.z.2* alters neural crest gene regulatory network, and decreases melanocyte and glial footprint. (A) Schematic design of microarray experiment. (B and C) Bubble plots representing gene ontology functions of differentially regulated genes with a log_2_ fold change ≤ −0.6 (downregulated, B) and ≥ 0.6 (upregulated, C) upon silencing Z2 in *sox10* +ve cells. (D) Heatmaps representing differential expression of key transcription factors in *sox10* +ve cells of Z2 MO as compared to control across melanocyte, glial and neuronal lineages. (E) WISH based expression pattern of melanocyte markers c*kit* and *dct* at 24hpf (F) Tg(*foxd3*:GFP) labelling of glial cells that mark undifferentiated glia and (bottom) WISH of glial markers *krox20* and *mbp* at 5dpf. (G) Tg(*nb*:dsRed) labelling of neurons in the spinal chord bundle (lateral view). (Bottom) WISH *isl2a* marker for cranial motor neurons. (H) Tg(*mitfa*:GFP) labelling of melanocytes, (bottom) WISH of *mitfa* at 24hpf. (I) Bright field images of control, Z2 MO and Z2 MO coinjected with *mitfa* mRNA at 48 hpf. (J) Bars represent mean ± SEM (n=3) of the number of head melanophores with at least 30 embryos each. (K) Schematic representation of the neural crest derived lineages, highlighting the dependence of melanocyte and glial cells on *h2a.z.2* Numbers in WISH images indicate frequency of the represented phenotype in total number of embryos analyzed. Scale bars represent 100 μm.

To verify the reduction in specific cell types, we adopted WISH for lineage specific differentiation markers and analysis of cell type specific transgenic lines that label NCC derivatives. *dct* and *kita* that label melanocytes, show decreased staining in the Z2 morphants (Fig 2E). Similar pattern of decreased staining was observed for differentiated schwann cells using markers *krox20* and *mbp* that labelled the lateral line associated glial cells (Fig 2F). Undifferentiated schwann cells, visualized by *Tg(foxD3:GFP),* showed substantial reduction. *Tg(nbt:DsRed)* marked enetric neurons remain unaffected (Supplementary Fig 2) and similarly WISH for *isl2a* showed no change (Fig 2G bottom panel). However, we observed the presence of ectopic neurons in the trunk, above the spinal chord neuron bundle, suggestive of a local increase in the neuronal population (Fig 2G). Labelling by *Tg(sox10:GFP)* as well as alcian blue staining of craniofacial appendages and fin cartilage showed no major changes, suggesting normal cartilage specification. However, we noticed a consistent decrease in jaw size as a result of defective ceratobranchial appendage spacing (Supplementary Fig 2). Together, our data suggests that *h2a.z.2* plays a selective role in the derivation of melanocytes and glial populations from the NCC. Interestingly, previous studies have shown that these two cell types emerge from a common progenitor (Raible and Eisen, 1994).

### *h2a.z.2* functions in conjunction with *mitf* to control melanocyte derivation

Micropthalmia associated transcription factor (*Mitf*), though seemingly dispensible for specification (Johnson et al., 2011), is the earliest known marker of lineage commitment and controls expression of downstream identity genes to enable melanocyte fate determination (Goding, 2000; Lister et al., 1999). We indeed observe a decrease in *mitfa* expression in sox10+ve cells, and this was further corroborated by WISH staining for *mitfa* (Fig 2H). Therefore we set out to decipher the interplay of *h2a.z.2* and *mitfa* in mediating melanocyte lineage commitment. A marked rescue in the melanocyte counts in *mitfa* mRNA injected Z2 morphants was noted (Fig I&J). The above data, confirms genetic interaction between *h2a.z.2* and *mitfa,* and *h2a.z.2* likely functions upstream of *mitfa* and hence could modulate melanocyte fate.

### Derivation of melanocyte from mouse embryonic stem cells is under the control of *H2a.z.2*

From these experiments it is likely that *H2a.z.2* plays a role in the specification/determination of an upstream progenitor population from which melanocytes arise. Additionally, decrease in glia in *h2a.z.2* morphant indicates that the bipotent progenitor of melanocytes and glial populations could be affected. Further, an increase in neurons is suggestive of a compensatory increase in neuronal progenitors. Our observations therefore indicate a heirarchical model of fate determination in the NCCs, which is in tune with recent studies (Le Douarin and Dupin, 2003; Raible and Eisen, 1994; Soldatov et al., 2019). We propose a model, wherein *h2a.z.2* would operate at the level of the tripotent glial/neuronal/melanocyte progenitor in specifying a melanocyte/glial fate (Fig 2K).

While the zebrafish based experiments helped us to narrow down the function of this variant in melanocyte derivation, cellular and molecular insights are required to establish its role in specification. To understand this molecular mechanistic link, we established a method to derive melanocytes from R1/E mouse embryonic stem cells (mESCs), wherein the mESCs were converted to melanocytes through an intermediate embryoid body state (EBs) (Fig 3A). Targeted gene expression analysis of NCC, melanocyte, and stemness related genes were performed at different days of melanocyte formation (Fig 3B). As anticipated, the expression of *Mitf* was induced at day 5 and further increased till day 16. Similarly, we observed the expression of pigmentation gene *Dct* to be induced during the course of experiment. We noticed that the expression of stemness factors *Oct4* and *Nanog* were drastically reduced during the course of induction, suggesting that the cells are undergoing differentiation. Strikingly, we noticed that the expression patterns of *H2a.z.2* and *H2a.z.1* were opposite to each other. While the expression of *H2a.z.2* was positively correlated with melanocyte specification, the expression of *H2a.z.1* was decreased. This pattern of *H2a.z.1* is similar to stemness factors *Oct4* and *Nanog*.

**Figure 3:**
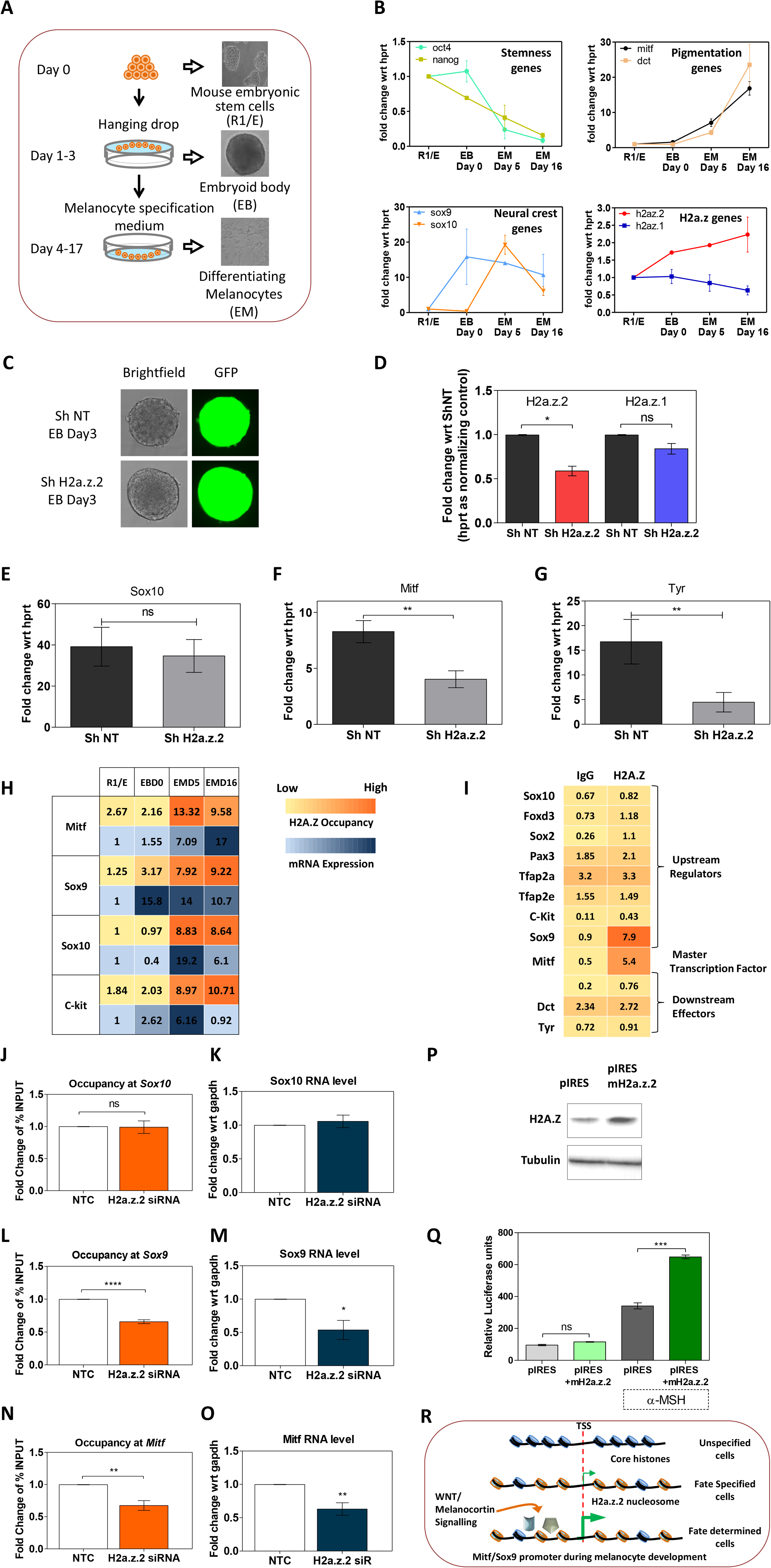
*H2.a.z.2* controls melanocyte derivation from mouse embryonic stem cells *in vitro*. (A) Schematic of melanocyte derivation from mouse R1/E stem cells. (B) Kinetics of expression patterns of stemness genes (*Oct4* and *Nanog*); pigmentation genes (*Mitf* and *Dct*); neural crest genes (*Sox9* and *Sox10*) along with *H2a.z.1* and *H2a.z.2* at different stages of melanocyte derivation from mouse R1/E stem cells. (C) Bright field and GFP images of embryoid bodies derived from R1/E cells silenced using lentiviral contruct for non-targetting or Z2 shRNA, encoding GFP marker. (D) Bar graphs representing qRT-PCR for *H2.a.z.1* (blue) and *H2.a.z.2* (red) in shZ2 cells compared to shNT at the embryoid body stage. (E-G) Bar graphs representing qRT-PCR analysis for *Sox10*, *Mitf* and *Tyr* (tyrosinase) on 6 days post induction of differentiation. (H) Heatmap representing the promoter occupancy of H2.A.Z (*H2.A.Z.1* and *H2.A.Z.2*) determined by Chromatin Immunoprecipitation (ChIP) and the corresponding mRNA levels of *Mitf*, *Sox9*, *Sox10* and *Ckit* during various stages of melanocyte derivation from R1/E cells. Numbers in the heat map represent percent enrichment of H2A.Z occupancy in ChIP and fold change compared to R1/E cells in the quantitative real-time PCR for the mRNA levels. (I) H2A.Z occupancy at the indicated promoters in melanocytes (depigmented B16 cells) is represented as a heat map of percent input for H2.A.Z as well as normal rabbit IgG. Numbers in the heat map represent percent enrichment of the promoter DNA in H2A.Z or IgG ChIP. (J,L,N) H2A.Z ChIP of *Sox10*, *Sox9* and *Mitf* promoters upon Z2 knockdown. (K,M,O) mRNA levels of *Sox10*, *Sox9* and *Mitf* upon Z2 knockdown. (P) Western blot for H2a.Z upon Z2 overexpression. (Q) Reporter assays for *Mitf* promoter by dual luciferase assay in control and H2.a.z.2 overexpressed cells in basal as well as α-Melanocyte Stimulating Hormone (α-MSH) treated B16 cells for 24h. Bar graphs represent mean±SEM (n=3). (R) Schematic representation of the role of *H2.a.z.2* in modulating *Mitf* promoter.

To illustrate the functional role of *H2a.z.2* in mESCs, we silenced *H2a.z.2* and studied melanocyte derivation. In the cells stably expressing short hairpin RNA that targets h2a.z.2 (shZ2), a reduction of around 50% in *H2a.z.2* mRNA levels could be observed, while *H2a.z.1* levels remained unaltered (Fig 3C). Interestingly, previous studies report that knockdown of *H2a.z.1* in embryonic stem cells leads to the formation of defective embryoid bodies with irregular structural integrity (Hu et al., 2013). We did not observe significant difference in the size and morphology of the *H2a.z.2* depleted EBs (Fig 3D), further strengthening a differential role for the two proteins, at least during early developmental stages.

Upon induction of melanocytes from R1/E cells stably expressing shZ2, compared to nontargetting shRNA (shNT), the expression of neural crest gene *Sox10* remained unaltered. This observation is strikingly similar to Z2 MO scenario in zebrafish embryos (Fig 3E). A significant decrease in the mRNA level of the master transcription factor *Mitf* and its target gene *Tyr* in shZ2 cells as compared to shNT cells was observed. Thus highlighting a conserved role for *H2a.z.2* in melanocyte specification across zebrafish and mouse. Having set up the cell based model of melanocyte specification and the effect of *H2a.z.2* in derivation of melanocytes in this model, we set out to identify effectors and the mechanism by which *H2a.z.2* could mediate melanocyte specification.

### H2A.Z.2 occupies *Sox9* and *Mitf* promoters, and controls *Mitf* inducibility

Both H2A.Z.1 and H2A.Z.2 occupy the nucleosomes in promoter as well as enhancer regions immediate to the transcription start site (TSS) of the downstream genes. This occupancy can either facilitate (Bargaje et al., 2012; Gevry et al., 2009; Zovkic et al., 2014) or repress (Dai et al., 2017; Hardy et al., 2009) transcription, in a context and a model system dependent manner. Partly, this discrepancy could be attributed to the near identical proteins that are indistinguishable in chromatin immunoprecipitation (ChIP) studies and to the complex nature of chromatin changes brought about by H2A.Z. We therefore employed a silencing strategy followed by ChIP to address the molecular mechanism selective to Z2.

H2A.Z ChIP followed by q-PCR for the promoter proximal nucleosomes of *Mitf*, and other critical upstream effectors *Sox9*, *Sox10* and *Ckit* was performed across various stages of melanocyte specification in the mESC model. We then compared the nucleosome occupancy of H2A.Z to their mRNA level changes at these stages of differentiation. All these four promoters showed increased occupancy during the course of melanocyte formation from mESCs. Interestingly, we observed that mRNA expression of *Mitf*, positively correlated with the occupancy of H2A.Z (H2A.Z.1+H2A.Z.2) in its promoter, suggesting that H2A.Z facilitates induction of *Mitf* expression possibly by modulating the accessibility of *Mitf* promoter (Fig 3H).

Though our progressive induction model provided a window to assess kinetic changes in promoter occupancy, heterogeneity in the cell populations arising during the course of melanocyte induction, limited our interpretations. Therefore, we employed a complementary approach wherein we utilized B16 melanoma cells that would represent specified melanocytes. ChIP with H2A.Z antibody was carried out in B16 cells to check the occupancy in promoter regions of genes related to *Mitf,* and upstream regulators as well as genes downstream to it in melanocytes. *Mitf* and *Sox9* promoters showed considerable enrichment of 5-7% over input (Fig 3I), suggesting that these genes might be under H2A.Z control. In contrast, several pigmentation genes like *Dct*, *Tyr*, *Tyrp1, Sox10* etc. were not enriched for H2A.Z in B16 cells. This is in tune with previous studies on H2A.Z in human melanoma cells (Vardabasso et al., 2015) and our meta-analysis further corroborated this observation (Supplementary Fig 3).

Knockdown of *H2a.z.2* in B16 cells with siRNA resulted in a consistent 70-80% decrease in mRNA levels of *H2a.z.2*, while *H2a.z.1* levels remained unchanged (Supplementary Fig 3). H2A.Z ChIP-qPCR was carried out for *Mitf*, *Sox9* and *Sox10* promoter regions in *H2a.Z.2* knockdown B16 cells (Fig 3J,L&N). We observed decreased occupancy of H2A.Z upon Z2 knockdown (Fig 3L&N) and a correlated decrease in mRNA expression of *Mitf* and *Sox9* (Fig 3M&O) suggested a direct consequence of Z2 incorporation. In contrast, *Sox10* promoter, which had minimal occupancy in B16 cells, showed no change upon *H2a.z.2* silencing. Correspondingly, the *Sox10* mRNA levels remained unaltered (Fig 3K). Promoter specific luciferase constructs, further demonstrated *Mitf* and *Sox9* promoters to be regulated by H2A.Z.2 (Supplementary Fig 3). Taking into account these observations we conclude that Z2 modulates the promoter region of MITF and facilitates its induction, the absence of which leads to decreased Mitf expression resulting in reduced number of specified melanocytes.

The effect of H2A.Z2 occupancy on *Mitf* promoter was addressed in reporter assay with the overexpression of *H2a.Z.2* in B16 cells. While there was no change in the *Mitf* promoter activity under basal conditions in Z2 overexpressing cells, MSH stimulation resulted in 7-fold upregulation, which was much higher than control empty vector transfected cells (Fig 3P&Q). We thus conclude that positioning of H2A.Z.2 upstream of *Mitf* facilitates its induction by appropriate cues. Occupancy of H2A.Z.2 renders chromatin state of the promoter to be highly responsive to activating signals in the form of transcription factors. The role of H2A.Z.2 delineated in the current study shows striking similarity with its role in brain, wherein incorporation of H2A.Z mediates dynamic gene expression changes associated with memory formation (Zovkic et al., 2014). It also is interesting to note that upon Z2 silencing in the zebrafish system as well, *sox9a* and *sox9b* show reduction in the NCC populations (Fig 2D). Given the upstream role of *Sox9* as a positive regulator of *Mitf*, it is likely that this may play an additional role in melanocyte specification via *Mitf*. Thereby we have identified a histone code upstream of the transcription factor network that specifies melanocytes from the pluripotent NCCs (Fig 3R).

### *H2a.z.2* is required for adult melanocyte regeneration

Targeted knockout of *H2a.z.1* has been attempted in several organisms and found to be embryonically lethal. However, the Z2 variant has not been not systematically studied. CRISPR-Cas9 based targeted deletion of *h2a.z.2* in zebrafish resulted in severe lethality and gross deformities (Supplementary Fig 4). Genotyping of *h2a.z.2* locus from these embryos revealed several in-dels resulting in large scale disruptions of this gene. Though *h2a.z.2* seems to be essential for early development, we could isolate three *h2a.z.2* mutant lines by targetting the coding region downstream to the core histone domain which map to the C-terminal extension region beyond αC domain (Fig 4A). This region is unstructured, and is involved in dynamic docking with histone H3/H4 during nucleosome formation (Bonisch et al., 2012). Extensive work based on systematic mutagenesis of *H2A.Z* (Wratting et al., 2012) and studies involving a primate specific, alternately spliced isoform *H2A.Z.2.2* (Bonisch et al., 2012), enabled prediction of nucleosome stability of the mutants. The truncation of this region would have least effect on nucleosome assembly, and replacement by a shorter stretch is expected to have a moderate effect. Further, elongation of the C-terminal extension is known to have severe effect based on yeast H2A.Z study (Wratting et al., 2012). Hence the nucleosome stability would be H2A.Z.2^Gln125fsX144^ < H2A.Z.2.2 = H2A.Z.2^Ile101fsX110^ < H2A.Z.2^Δ100-120^ < H2A.Z.2 (Fig 4A, right). We observe pigmentation phenotype severity to be in tune with the expected effect of mutations (Fig 4B).

**Figure 4:**
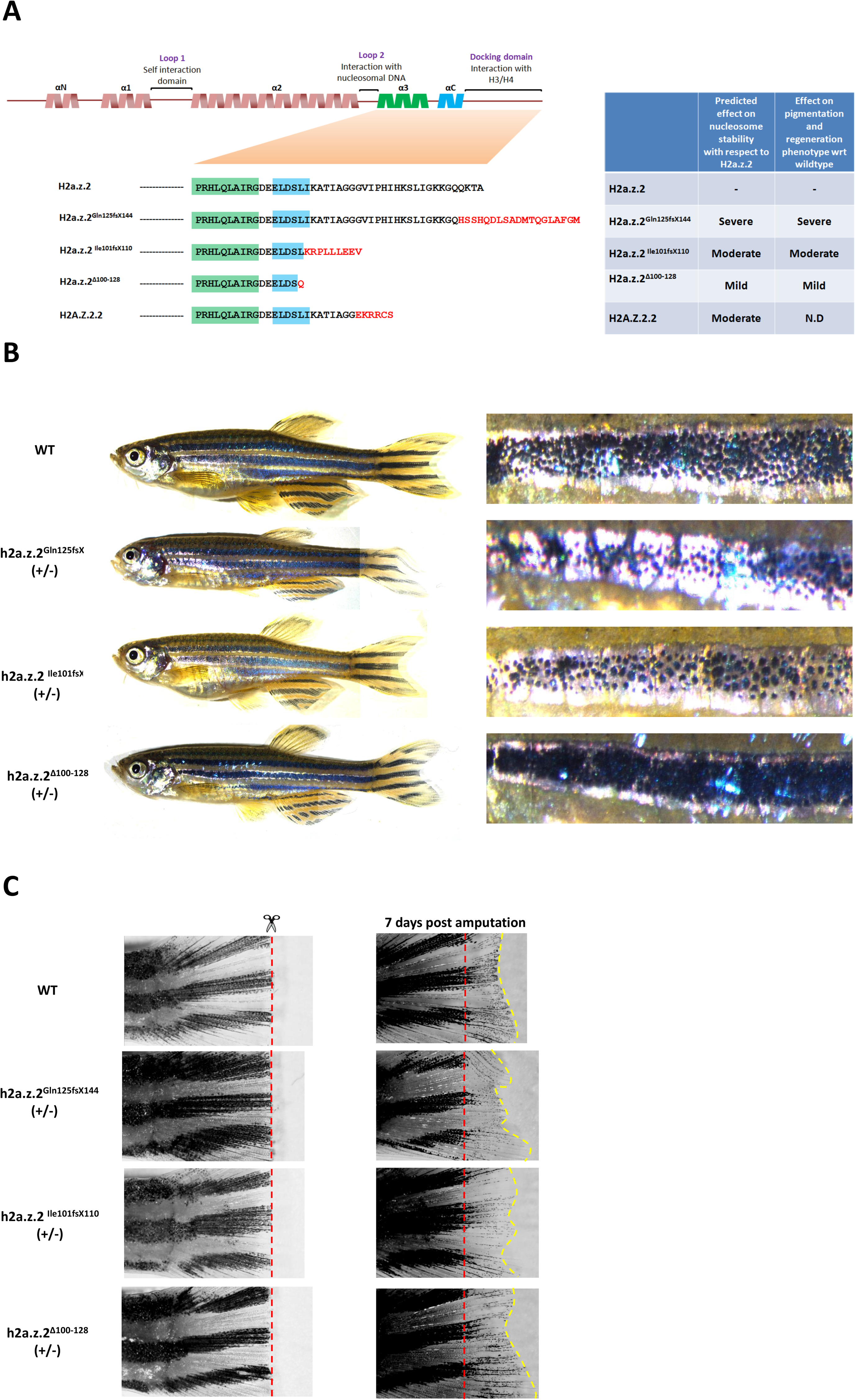
Targeted mutation of *h2a.z.2* affects melanocyte specification and regeneration. (A) Orthology based schematic representation of the secondary structure of zebrafish H2A.Z.2. The functional elements in the protein are denoted above.(below) Sequence of the amino acids in H2A.Z.2, its variant H2A.Z.2.2 and the three CRISPR mutants generated in this study. (B) Representative images of wild type and Z2 mutant adult fishes. (right) zoomed in images of first ventral stripe. (Adult fishes were imaged in parts and stitched) (D) Fin clip and regeneration images of adult wild type and Z2 mutant animals. Right panel represents fins after 7 days of regeneration.

Interestingly, in the mutant heterozygous lines we observed less melanocytes, confirming partial loss of function and a dominant nature of the mutations. In the adult animals, role of H2A.Z.2 in specification could be verified by studying melanocyte regeneration, a process wherein differentiated melanocytes arise from a pre-specified progenitor pool and could be experimentally studied using finclip experiments (Hultman et al., 2009). Herein the adult zebrafish fin is cut and allowed to regenerate for a week, during which the quiescent progenitors are stimulated and undergo the process of differentiation to generate pigmented cells. We noticed that all the three *h2a.z.2* mutants demonstrated comparable fin growth, however they displayed decreased melanocyte regeneration, in tune with the severity of the ventral stripe pigmentation observed in these mutants (Fig 4C). The NCC-derived fate specified progenitors establish in skin, and respond to melanocyte loss by providing requisite melanocytes. Decreased recall of melanocytes indicates that the progenitors are inadequately populated, and endorses the role of *h2a.z.2* in melanocyte fate specification. Thereby using targetted genome engineering, we unequivocally demonstrate the role of H2A.Z.2 in establishing melanocyte progenitor population, evident not just at the embryonic stage but also during adult melanocyte regeneration.

## Discussion

Decoding cell fate decision has emerged as an intense area of research, primarily due the immense application potential it has in futuristic regenerative interventions. Fate establishment is perceived to be a continuum, wherein progressive fate bias culminates in determination and is controlled by appropriate external signals. Currently, we understand deterministic events better, due to the characteristic gene expression signatures orchestrated by key transcription factors. However, the underlying molecular mechanisms behind fate-biasing specification events remains unexplored. Non-deterministic nature of fate bias induced during specification necessitates a modulatory role for the specifier and epigenetic factors are perfectly suited. Histone modifications and deposition of variants could function in conjunction with transcription factors and offer a greater plasticity to modulate the cell fate. In this study, we unequivocally establish the role of H2A.Z.2 as a rheostat for *Mitf* induction by facilitating chromatin accessibility to upstream activating signals and propose a model for melanocyte lineage specification, wherein H2A.Z.2 occupancy in *Mitf* acts as a key specifying signal.

Three of the fate-biasing transcriptional programs in neural crest lineage are involved in suppressing the melanocyte fate. These include *Sox2* (Adameyko et al., 2012) and *Neurog2* (Soldatov et al., 2019) from neural, and *FoxD3* (Johnson et al., 2011) from the glial lineage, all of which actively suppress *Mitf* expression. *H2a.z.2* in contrast is a positive enabler of melanocyte fate bias, as its overexpression facilitates *Mitf* induction. Heirarchical fate biasing model necessitates such opposing programs to ensure defined populations of emerging, differentiated cells. The observed concomitant decrease in the glial population suggests that H2A.Z.2 could operate as a specifier at the tripotent melanocyte/glial/neural progenitor level, driving the allocation towards a melanocyte-glial bipotent fate. *Sox9* regulation by H2a.z.2 could have a dual role of promoting *Mitf* expression in melanocytes and also facilitating the glial specific programs (Cheung and Briscoe, 2003).

Regenerative melanocytes arise from NCC-derived stem cells that are positioned along the lateral line in close juxtaposition with dorsal root ganglia in zebrafish embryos (Dooley et al., 2013). In the adult zebrafish, precursor cells directly differentiate into mature melanocytes and also divide to yield additional lineage-restricted cells (Iyengar et al., 2015). In mammalian skin, this population resembles nerve associated melanocyte precursors (Adameyko et al., 2009) and the hair follicle bulge region progenitors, that give rise to melanocytes at every hair cycle (Nishimura et al., 2002). These progenitors are thought to restore epidermal pigmentation after wound healing and during melanocyte regeneration in conditions such as vitiligo (Chou et al., 2013).

Interestingly, though the establishment of melanocyte stem cells is seemingly independent of *Mitf* (Johnson et al., 2011), all the downstream events are orchestrated by this master regulator. Given this central role of *Mitf* in fate determination, the “melanocyte specification factor” is likely to function upstream of it. The transient epigenetic memory conferred by the incorporation of H2A.Z.2 at *Mitf* promoter would not just facilitate its expression upon external cues such as WNT ligands, it would also enable retention of this memory in transiently amplifying cells. The *Mitf*-inducing Wnt signaling pathway triggers melanocyte regeneration possibly by activating this poised state of *Mitf* promoter. Thereby such a histone code elucidated in our study enables retention of plasticity, a hallmark of the specified state. Identification of such upstream codes would enable futuristic regenerative therapeutic approaches from induced pluripotent cells, by employing selective activation of lineage determinants.

## Materials and Methods

### Ethics Statement

Fish experiments were performed in strict accordance with the institutional animal ethics approval (IAEC) of the CSIR-Institute of Genomics and Integrative Biology, India (Proposal No 45a). All efforts were made to minimize animal suffering.

### Zebrafish lines and Maintenance

Zebrafish line ASWT were bred, raised and maintained at 28.5 ^0^C according to standard protocols (Westerfield, 2000) and were housed at the CSIR-Institute of Genomics and Integrative Biology (IGIB), Mathura Road New Delhi, India. Embryos were staged both using timing (hours post fertilization (hpf); days post fertilization (dpf)) and morphological features according to (Kimmel et al., 1995). Embryos older than 24hpf were treated with 0.003% PTU (1-phenyl-2-thiourea) to inhibit pigment formation, aiding fluorescent imaging and RNA *in situ* hybridization analysis, as and when the experimentation demanded depigmenting the animals. *ftyrp4*:GFP plasmid was a kind gift from Dr. Xiangyun Wei (University of Pennsylvania) (Zou et al., 2006) and the construct was injected at a concentration of 10-20 pg alongwith 50-75 pg of Tol2 transposase mRNA into one cell stage zebrafish embryos to create the transgenic line at CSIR-IGIB. Founder lines were established and propagated, which showed expression pattern similar to pt101 as reported in (Zou et al., 2006). The details of zebrafish lines used in this study are provided in key resources table.

### Morpholino injections

Morpholinos for blocking translational initiation or splicing of RNA were designed by GeneTools^®^ (Supplementary table 1). Minimum effective concentration was determined for each morpholino (MOs) using dosage titration experiments. Concentrations of various MOs used in this study are as follows: 3 – 3.5 ng of *h2a.z.2* translation block (Z2 MO), 4 – 4.5 ng of *h2a.z.2* splice block (Z2 SB MO), 2 ng of *h2a.z.1* splice block (Z1 MO) (Sivasubbu et al., 2006) and 0.8ng of double targeting (*h2a.z.1* and *h2a.z.2*) translational block (Z1 + Z2 MO). Standard control morpholino from gene tools was used as a control and dosed accordingly. p53 morpholino was co-injected at 1:1 ratio with corresponding gene targeting morpholinos.

### Capped RNA injections

For rescue experiments, mouse *H2a.z.1* (Z1-RNA) and *H2a.z.2* (Z2-RNA) gene were amplified using primers as listed in (Related to STAR methods: Oligonucleotides). Similarly, the zebrafish *mitfa* gene was amplified using PCR (Related to STAR methods: Oligonucleotides) and cloned into Zero Blunt TOPO vector (Thermo Scientific^®^; K287540) according to manufacturer protocols. Capped and PolyA tail RNA was synthesized by *in vitro* transcription using T7-Ultra mRNA synthesis kit (Thermo scientific^®^; AM1345) followed by polyA tailing according to manufacturer’s protocol. 100pg (for *H2a.z.1* and *H2a.z.2*) or 50pg (for *mitfa*) was co-injected into 1-cell stage embryos along with the Z2 morpholino.

### Whole mount RNA *in situ* hybridization

RNA *in situ* hybridization was performed as described in (Thisse and Thisse, 2008). Riboprobes of various genes foxd3, sox10, crestin, krox20, sox9a (Stewart et al., 2006), mbp (Brosamle and Halpern, 2002), neuroD (van der Velden et al., 2013), isl2a (Appel et al., 1995), sox9b (Yan et al., 2005), tfap2a, mitfa, c-kit, dct (Van Otterloo et al., 2010) were used for this study. Zebrafish *h2a.z.2* exon5 – 3’UTR (307 bp) was amplified using primers listed in (Related to STAR methods: Oligonucleotides). The product was subsequently cloned into pCR4-TOPO (Thermo Scientific^®^; K287540) and was used as template for *in vitro* transcription of antisense probe using Megascript T7 (Thermo Scientific^®^, AM1334).

### Imaging

Bright field imaging for live imaging and gene expression pattern (WISH) studies of zebrafish embryos was performed using Zeiss (Stemi 2000C). Zeiss axioscope A1 microscope (with AxiocamHRc) was used for fluorescence imaging. Zeiss proprietary software was used to capture images and processed and analyzed in Adobe Photoshop CS3.

### Imaging flow cytometry based population analysis

Cell counts for sox10:EGFP, mitfa:GFP, ftyrp1:GFP positive cells and ASWT embryos (for melanophore count) were performed using AMNIS^®^ Imaging FACS. Briefly, Embryos were dechorinated using pronase (5mg/ml) (Sigma, P8811) for 10-15 mins and collected in microfuge tubes. The embryos were deyolked in ice cold ringer’s solution using micropipette tip and spun at 100g for 2 minutes at cooled table top centrifuge at 4^0^C (Eppendorf^®^ centrifuge 5418R). The supernatant was discarded and the embryo bodies were trypsinized using TrypLE Express (Thermoscientific^®^, 12604039) for 15 or 30 minutes for 24hpf or 48 hpf embryos respectively at room temperature. The cell suspension was passed through 70μm cell strainer and washed twice with ice cold phosphate buffered saline. The cell suspension was analyzed in the Imaging Flow cytometry system and at least 50,000 – 1,00,000 images were captured and processed using IDEAS (AMNIS^®^) software.

### Morphant *Tg(sox10: GFP)* cell sorting and microarray analysis

Control MO and Z2 MO were injected in 1 cell stage *Tg(sox10:GFP)* embryos and allowed to develop till 15 hpf. The embryos were processed in a similar manner as that for Imaging flow cytometry analysis. Finally, the trypsinised cells were resuspended in ice cold PBS + 10% Fetal bovine serum and subjected to FACS (BD Bioscience FACSAria^TM^ III). The GFP positive cells were sorted and collected in RP1 buffer and RNA was isolated using Nucleospin RNA XS kit (Macherey Nagel, 740902) according to manufacturer’s protocol. Microarray and analysis was outsourced to Genotypic Technology (Bengaluru, India).

### Cell line maintenance and culturing conditions

B16 mouse melanoma cells were cultured in DMEM-High glucose media (Sigma; D5648) supplemented with 10% FBS (Thermo Scientific^®^; 10270-106) at 5% CO2 (Eppendorf^®^ New Brunswick Galaxy170S). Media was changed every day and cells were passaged upon reaching 80% confluence. Mouse R1 embryonic stem cells (R1/E) were cultured in DMEM Glutamax media supplemented with sodium pyruvate (Thermo Scientific^®^; 10569-010), MEM-NEAA (Thermo Scientific^®^; 11140-050), Beta-mercaptoethanol (Thermo Scientific^®^; 31350-010) and 20% PANSERA (Pan Biotech; P30-2602) at 5% CO2. Media was changed every day and cells were passaged every other day.

### Melanocyte generation from R1/E mouse embryonic stem cells

Melanocytes were generated from R1/E mESCs using modified protocol from (Yang et al., 2011). For embryoid body formation, cells were trypsinsed and 50,000 cells/ml suspensions was prepared. 10ul of this suspension was placed in the lid of a 90 mm petridish and inverted upon the base of the petriplate filled with 15 ml of autoclaved MilliQ. After 4 days the embryoid bodies were transferred to 24 well or 6 well plates coated with collagen (Thermo Scientific^®^; A10483-01) containing Melanocyte conversion media (45% DMEM – High glucose, 45% Reconstituted M254 (Thermo Scientific^®^; M254CF), 10% FBS, 50ng/ml SCF (Peprotech, 300-07-10), 100nM Endothelin-3 (Sigma, E9137), 20pM Cholera toxin (Sigma, C8052), 0.5uM Dexamethasone (Sigma, D1756), 50ng/ml WNT3a (Peprotech; 315-20-10), 4ng/ml beta-FGF (Thermo Scientific^®^; RFGFB50), 50nM PMA (Sigma, P1585),1x N2 Supplement (Thermo Scientific^®^; 17502-048) and 1X Anti Anti (Thermo Scientific^®^; 15240-062). Cells were collected at specific time points for downstream analysis.

### siRNA and plasmid transfections

2 x 10^5^ B16 melanoma cells were seeded in 6 well plates prior to the day of transfection. Next day 100nM of non targeting control siRNA, *H2a.z.2* siRNA (siRNA pool of 5 siRNAs) (Dharmacon™ ON-TARGETplus; L-063612-01-0005) or *H2a.z.1* siRNA (siRNA pool of 5 siRNAs) (Dharmacon™ ON-TARGETplus; L-042994-01-0005) was transfected according to manufacturer’s protocol using Dharmafect (Dharmacon^TM^, T-2001) transfection reagent. Cells were collected at specified time points for downstream analysis. Plasmid transfections were performed using Lipofectamine 2000 (Thermo Scientific^®^; 11668019)

### RNA isolation, cDNA synthesis and quantitative Real time PCR

RNA was isolated from zebrafish embryos or B16 melanoma cells or R1/E cells using Nucleospin Triprep (Macherey Nagel, 740966) according to manufacturer’s protocols. cDNA was synthesized using Superscript III First strand cDNA synthesis kit (Thermo Scientific^®^; 1800051). Quantitative real time PCR was carried out as described (Pfaffl, 2001) on ROCHE Lightcycler^®^ Real time PCR system. For quantification, the relative standard curve method was used (as described by the manufacturer) to generate raw values representing arbitrary units of RNA transcripts. Data analysis was performed using 2^-ΔΔCT^ method.

### shRNA transduction in R1/E ESCs

Lentiviral packaging and transduction was performed as reported previously by (Motiani et al, 2013a, b). Briefly, lentiviral packaging was done by co-transfecting pVSVG, pdR8.2, and *H2a.z.2* shRNA or Non-targeting shRNA in HEK cells. Two days post-transfections, lentiviral particles were collected from cell supernatant. 1:100 HEPES pH 7.25 1M and 1:100 Polybrene 0.4 mg/ml was added to lentiviral supernatant. Embryonic stem cell media (ESCM) and lentiviral cocktail were mixed in 1:1 ratio and added to R1/E ESCs. The occulate was spun for 30 min at 37^0^ C and 2500rpm. The cells were incubated at 37^0^ C for 10-12 hours and then media was changed to fresh ESCM. Cells were checked for GFP fluorescence at 48 hours after transduction in R1/E cells and were selected using 0.5μg/ml puromycin for a week. Experiments were performed with FACS sorted GFP +ve cells propagated as a pool.

### Chromatin Immunoprecipitation and qPCR

ChIP assays were performed according to protocol provided by Upstate Biotechnology with modifications as suggested in Fast ChIP protocol. ChIP assays were performed using anti-H2A.Z antibody (abcam; ab4174). Anti-Rabbit IgG (Thermo Scientific®; 02-6102) was used for isotype control in all the cell lines. Briefly, B16 Melanoma cells or R1/E ESCs were fixed with 10% formalin (Sigma; HT501128) and incubated at 37^0^C for 10 minutes. 2.5 M Glycine (Sigma; 50046) was added to the cells and again incubated at 37^0^ C for 10 minutes. Cells were washed with ice cold 1X PBS containing protease inhibitors. Cells were then scraped and centrifuged at 1000 rpm for 5 minutes at 4^0^C. The cell pellet was lysed in SDS lysis buffer (1% SDS (Sigma; L3771), 10mM EDTA (Sigma; E6758), 50mM TRIS (Sigma; T6066) (pH 8.1)) on ice for 30 minutes. The cells were then sonicated on bioruptor (DIAGENODE) in ice. The chromatin lysate was then estimated for protein content using BCA kit (Thermo Scientific^®^; 23225). The samples were de-crosslinked overnight at 65^0^C, and deproteinised using with Proteinase K (Sigma; P4850). Sheared DNA was column purified using PCR purification kit (QIAGEN; 28104) and estimated using QUBIT dsDNA HS kit (Thermo Scientific^®^; Q32851). ChIP was performed using 3-5 μg of the respective antibody added to equal amount of chromatin across samples and incubated overnight at 4^0^C. 10% of chromatin was kept separately as input. Next day, 100ul of Protein A agarose beads (G-Bioscience, 786-283) were added to the mixture of chromatin and antibody and incubated for 4-6 hours at 4^0^C. After incubation the beads were washed twice with Low salt Buffer (0.1 % SDS, 1% Triton X 100 (Sigma; T8787), 2mM EDTA, 20mM Tris HCl (pH 8), 150mM NaCl (Sigma; S3014)), high salt buffer (0.1 % SDS, 1% Triton X 100, 2mM EDTA, 20mM Tris HCl (pH 8), 500mM NaCl) and LiCl buffer (0.25 M LiCl (Sigma; 62476), 1% Igepal CA-630 (Sigma; I8896), 1mM EDTA, 10mM Tris HCl (pH 8), 1% deoxycholate (Sigma; D6750)). Finally the beads were washed with Tris EDTA buffer. The beads were then incubated with 100μl elution buffer (1% SDS, 0.75% sodium bicarbonate (Sigma; S5761)) and 1 μl of 20mg/ml proteinase K for 15 minutes; elution was performed again with 100μl for another 15 minutes. Subsequently the samples were kept for 12 – 14 h at 65^0^C for reverse crosslinking. After incubation, the samples were column purified using PCR purification kit; the inputs from each of the samples were also included in the purification step. SYBR green (KAPABiosystems; KK4601) based qRT PCR was setup using eluted DNA and graphs were plotted as percentage input. Primer sequence for target genes are provided in Supplementary Table under Oligonucleotide sequences).

### Luciferase assay

Mouse *Mitf*-luciferase (Kind gift from Dr Krishnamurthy Natarajan, Jawaharal Nehru University) and *Sox9*-luciferase (Kind gift from Dr Peter Koopman, The University of Queensland) constructs were co-transfected (at ratio of 1:10) with renilla luciferase (pRL-TK vector, PROMEGA) in Non Targeting Control or *H2a.z.2* siRNA treated B16 melanoma cells. After 24 hr the cells were lysed with passive lysis buffer and luciferase assays were performed according to manufacturer’s protocols (Dual luciferase reporter assay system; Promega). Alpha MSH (Sigma; M4135) treatment was provided 6 hours post transfection of luciferase constructs.

### CRISPR based mutagenesis

sgRNA targeting zebrafish *h2a.z.2* gene were selected from ECRISP (http://www.e-crisp.org/E-CRISP/) online tool using default parameters. Primers were designed for generating the complete sgRNA (Related to STAR methods: Oligonucleotides) using annealing PCR. In vitro transcription was performed on these PCR products using T7 Megashortscript kit (Thermo Scientific^®^; AM1354) according to manufacturer’s protocols. 100pg of sgRNA was injected with 500 pg of spCAS9 protein (kind gift from Dr Debojyoti Chakraborty, CSIR-IGIB, India). The F0 embryos were grown to adulthood and then were out crossed with ASWT to give rise to F1 animals. In F1 generation, genomic DNA was isolated from fin clips of putative mutants and the target region was amplified using primers provided in (Related to STAR methods: Oligonucleotides). The PCR products were subjected to Sanger sequencing for confirming mutations.

We used the F2 generation h2a.z.2 mutants namely, h2a.z.2^Gln125fsX144^, h2a.z.2^Ile101fsX110^ and h2a.z.2^Δ100-128^ for this study. In the H2a.z.2^Gln125fsX144^ mutants we observe 10 base deletion in exon5, which leads to skipping of the stop codon and adds an additional 14 amino acids to the H2a.z.2 protein sequence. The h2a.z.2^Ile101fsX110^ mutants are characterized by a 3 bp deletion leading to change in amino acid sequences from 100-109 and generating a premature stop codon. The h2a.z.2^Δ100-128^ mutants are characterized by 6 base deletion and 3 base insertion leading to premature stop codon at the 100^th^ amino acid.

### Melanocyte regeneration via fin amputation

Fin amputation experiments were performed on heterozygous F1 mutant animals. Mature zebrafish adult (5-10 months) were anesthetized and the distal two-thirds of the caudal fin amputated with a scalpel. The amputated fin was imaged immediately in a stereozoom microscope (Zeiss Stemi 2000C). The fish were then returned to fresh water at 25°C on a regular feeding schedule for the duration of the experiment. The fin is allowed to regenerate for a week, during which the quiescent MSCs are stimulated and undergo the process of differentiation to generate pigmented cells. The tail fin of the fish was imaged again after 7 days and melanocyte regeneration was assessed.

### Statistical analysis and Graphs

Student’s t test was performed to obtain statistical significance in the data. Asterisk on the error bar corresponds to *(P ≤ 0.05), ** (P ≤ 0.01), *** (P ≤ 0.001), **** (P ≤ 0.0001) and ns (P > 0.05). Graphs were plotted using Graphpad prism.

## Acknowledgements

This work was supported by the Council for Scientific and Industrial Research (CSIR), India through grant (TOUCH-BSC0302, GRAFT-MLP1810 and OLP1118) and Department of Biotechnology through the grant (GAP0182). We acknowledge Dr Chetana Sachidanandan (CSIR-IGIB, New Delhi, India) for providing several zebrafish lines and WISH probes. We acknowledge Dr Aswini Babu for assistance with WISH experiments and Tg (sox10:GFP) embryos FACS sorting. We acknowledge the Imaging and FACS facility of CSIR-IGIB. DAR acknowledges senior research fellowship from ICMR, India.

## Author contributions

D.A.R and T.N.V designed experiments. D.A.R and Y.J.S performed experiments pertaining to zebrafish, D.A.R, Y.J.S, V.G and J.T performed experiments with cultured cells. D.A.R, S.S, R.S.G, R.M and T.N.V were involved in the design and execution of zebrafish experiments. D.A.R, S.S, R.S.G and T.N.V were involved in data analysis, interpretations and writing of the manuscript.

## Data Availability

Microarray data of isolated sox10 positive cells is deposited as NCBI GEO data set GSE133141

## Expanded View Figure Legends

**Suppl Fig 1: Validating H2a.z.2 specific pigmentation phenotype using multiple silencing approaches**

(A) Schematic representing the target region for H2a.z.2 MO (Z.2), H2a.z.1 MO (Z.1) and Z.2 & Z.1 MO.

(B) Brightfield images representing 48 hpf embryos injected with Control MO, Z.2 MO, Z.1 MO, Z.1& Z.2 MO.

(C) Brightfield images showing dorsal and lateral view of Control MO, Z.2 MO and Z.2 SB MO embryos at 48 hpf.

(D) H2a.z.2 RT-PCR amplicons from control and Z.2 SB MO injected embryos depicting the mis-spliced product.

(E) Bar graphs representing relative p53 mRNA levels in Z.2 MO as compared to control MO.

(F) Grouped bar plots representing percentage survival and pigmentation phenotype observed across control MO, Z2 MO and Z.2 + p53 MO.

**Suppl Fig 2: Status of neural crest derivatives in H2a.z.2 morphants**

(A) Table enumerating the number of enteric, and trunk ectopic neurons observed in control and Z.2 morphants. Numbers represent mean ± SD counted manually across ~50 embryos.

(B) Fluorescence images of enteric neurons in control and Z.2 morphants.

(C) Alcian blue staining highlighting the craniofacial cartilage and fin cartilage in control and Z.2 morphants.

(D) Fluorescence images of *Tg(sox10:EGFP)* head region representing the craniofacial cartilage system in control and Z.2 morphants.

**Suppl Fig 3: H2A.Z occupancy and gene expression changes.**

(A-C) Metanalysis of chromatin immunoprecipitaton (ChIP) data in melanocyte derived lines from GSE68223.

(A) H2A.Z (Z.1 + Z.2) occupancy near transcription start site (TSS) in primary human melanocytes.

(B) H2A.Z.2-GFP occupancy near TSS in SK-Mel 147 metastatic melanoma cells.

(C) H2A.Z.1-GFP occupancy near TSS in SK-Mel 147 metastatic melanoma cells. Dotted line indicates the TSS. Gene names are displayed on right corresponding to their ChIP seq profiles. Percent enrichment relative to input in the ChIP-seq data is displayed to the left.

(D) Bar graph representing mRNA levels of *H2a.z.1* and *H2a.z.2* upon Z*.1* silencing (mean±SEM, N=3).

(E) Bar graph representing mRNA levels of *H2a.z.1* and *H2a.z.2* upon *Z.2* silencing (mean±SEM, N=3).

(F) Bar graphs representing the relative Sox9 luciferase activity of cells treated with non targeting control and Z.2 siRNA (mean±SEM, N=3).

(G) Bar plot representing relative mRNA levels of pigmentation related genes upon Z.2 and Z.1 silencing in B16 cells (mean±SEM, N=2).

**Suppl Fig 4: Targeted knockout of *h2a.z.2* is lethal in zebrafish**

(A) Schematic representing the region targeted by *h2a.z.2* sgRNA1.

(B) Sequences displaying representative mutations occurred upon *h2a.z.2* sgRNA1 injections leading to embryonic lethality; predicted protein sequences are displayed on the right hand side.

(C) Bright field images of SpCas9 and SpCas9 + sgRNA1 injected embryos at 6 and 36 hpf.

